# AI4AMP: Sequence-based antimicrobial peptides predictor using physicochemical properties-based encoding method and deep learning

**DOI:** 10.1101/2020.12.17.423359

**Authors:** Tzu-Tang Lin, Li-Yen Yang, I-Hsuan Lu, Wen-Chih Cheng, Zhe-Ren Hsu, Shu-Hwa Chen, Chung-Yen Lin

**Affiliations:** Institute of Information Science, Academia Sinica, TAIWAN

## Abstract

**Motivation:** Antimicrobial peptides (AMPs) are innate immune components that have aroused a great deal of interest among drug developers recently, as they may become a substitution for antibiotics. However, AMPs discovery through traditional wet-lab research is expensive and inefficient. Thus, we developed AI4AMP, a user-friendly web-server that provides an accurate prediction of the antimicrobial activity of a given protein sequence, to accelerate the process of AMP discovery.

**Results:** Our results show that our prediction model is superior to the existing AMP predictors.

**Availability:** AI4AMP is freely accessible at http://symbiosis.iis.sinica.edu.tw/PC_6/

## 1 Introduction

The growing resistance of bacteria to antibiotics is a massive threat to public health. Antimicrobial peptides are promising alternative candidates for antibiotics. However, the cost and effort of antimicrobial activity evaluation are formidable, resulting in the difficulty of screening useful AMPs. Therefore, we built AI4AMP website service, which helps to predict whether a peptide possesses antimicrobial activity, based on a deep learning model. Our AMP prediction model is based on two parts: AMP encoding and machine learning. The purpose of AMP encoding is to produce machine-readable data from a given protein sequence. In this paper, we proposed PC6, a novel protein-encoding method, providing the information of six physicochemical properties of each amino acid and the amino acid order in a protein sequence. It was suggested that, compared to traditional machine learning methods, deep learning has a higher capacity for dealing with complex data and learning useful information (LeCun *et al.*, 2015). In recent years, deep learning techniques have been applied to solve various kinds of protein prediction tasks, such as AMPs, protein-protein interaction (Sun *et al.,* 2017), and human leukocyte antigen complex (Vang and Xie, 2017). Those deep-learning models achieved outstanding performance, and therefore we adopted deep learning methods for our prediction model in AI4AMP. To sum up, AI4AMP is a website service that predicts sequences’ antimicrobial activity by implementing the PC6 encoding method and deep learning approach.

## 2 Materials and methods

### 2.1 Implementation and Workflow

AI4AMP is a website service that allows users to predict whether the interested peptide sequences are AMPs. The input protein sequences data should be in FASTA format, and AI4AMP would provide an output of a CSV file that contains prediction scores (ranging from 0 to 1) and prediction results (YES or NO) of each input protein sequence. The prediction score represents the probability that the input peptide is an AMP. Prediction results, a binary column stating whether or not the input protein sequence is an AMP, is based on the prediction score with a 0.5 threshold. Note that our prediction model only accepts input peptides constituted by 20 essential amino acids. If the input sequences contain unusual amino acids such as B, Z, U, and X, they will not be recognized and proceeded through our prediction model. If an input sequence is longer than 200 amino acids, a 200 amino acid long sliding window will cut the sequence into multiple sequences by moving 100 amino acids at a time; those multiple sequences will then proceed through our model independently. The workflow of AI4AMP is presented in **Figure 1**.

**Figure 1.**
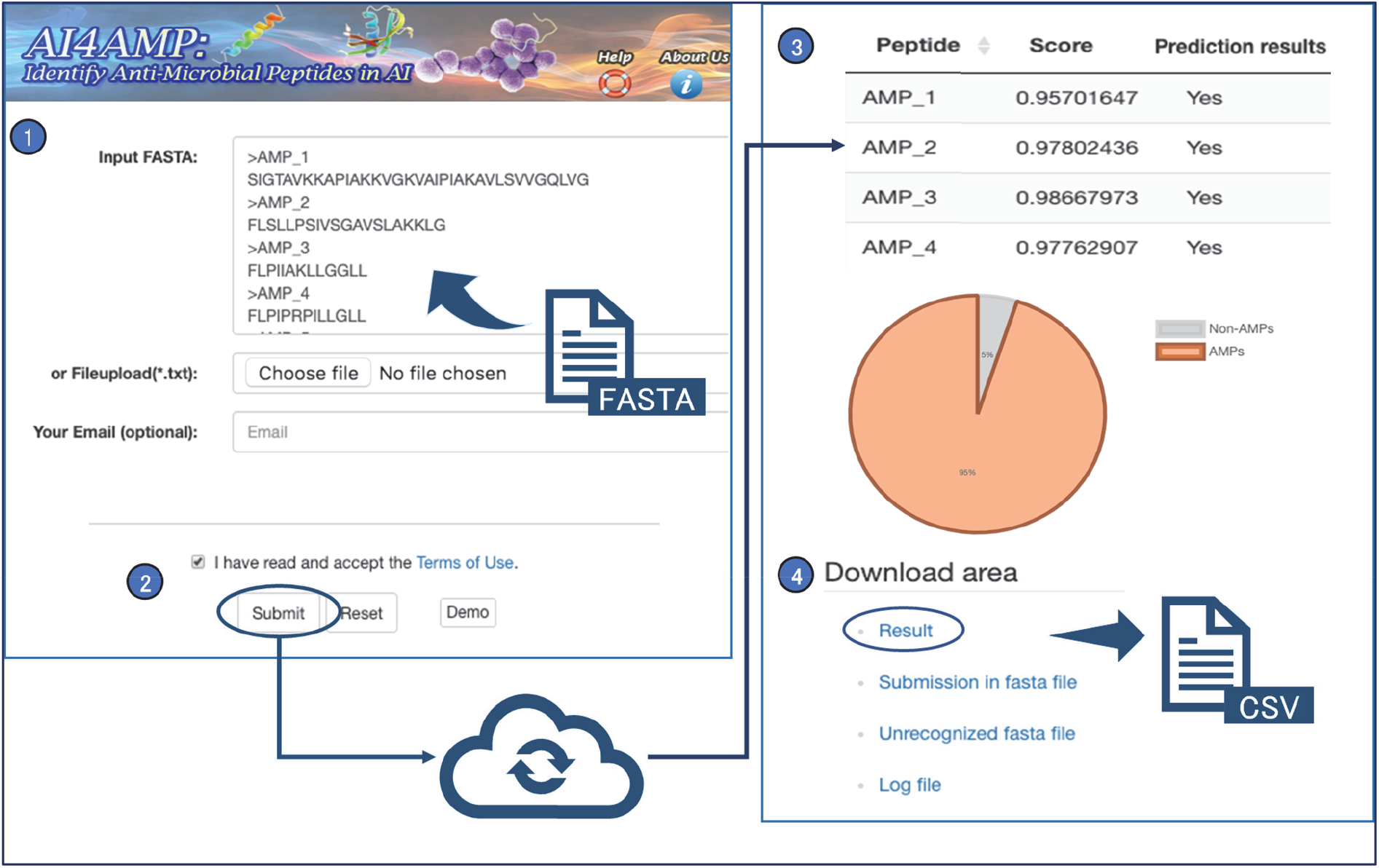
AI4AMP web server workflow. (1)(2) Users may either paste FASTA-format text in the “input FASTA” field or upload a FASTA file to “Fileupload(*txt).” Once the FASTA data is filled or uploaded, users may leave a valid email address in “Your Email” and click the “submit” button. (Results will be sent to user’s email only if the user fills out “Your Email.”) (3) In the result page, there will be a table demonstrating the prediction scores and prediction results of each peptide. There will also be a pie chart visualizing the proportion of AMP and non-AMP sequences of the input sequences. The total count, AMPs count, and non-AMPs count of the input sequences will also be shown underneath the pie chart. (4) Prediction result may be downloaded as a CSV file in the “Download area.”

### 2.2 Datasets

Our positive data were collected from four existing AMP databases: APD3, LAMP, CAMP3, and DRAMP. Our negative data were consisted of randomly generated sequences, non-AMP peptides from uniport, and negative data from an existing dataset (Veltri et al., 2018). For model tuning, we used 7056 data (the positive data were from APD3, CAMP3, and DRAMP), which were divided into 5715 training dataset, 635 validation dataset, and 706 testing dataset by splitting the data twice with 90:10 ratio. For external testing, 1130 data were obtained (the positive data were from the LAMP database only). Finally, we trained our model with all available data (13,246 data), producing the final prediction model of AI4AMP. (The detailed process is documented in Supplementary 1.)

### 2.3 Protein encoding methods

In our study, we developed a novel protein-encoding method, named PC6 encoding, which encodes protein sequences with the consideration of both the order and the physicochemical properties of amino acids of a given sequence. Our examination shows that PC6 encoding performs the best among the four methods. Therefore, we adopted PC6 encoding in AI4AMP. (Detailed PC6 encoding method may be found in Supplement 2.)

### 2.4 Deep learning

We implemented Keras, a high-level neural networks API, in our neural network. Grid search was performed to adjust hyperparameters such as learning rate, batch size, optimizer. (Model architecture and hyperparameter settings of AI4AMP are demonstrated in Supplementary 3.)

## 3 Result

AI4AMP is a user-friendly website service that demonstrated high accuracy of AMP prediction. We compared our model performance with the other state-of-the-art predictors: Antimicrobial Peptide Scanner vr.2 (APS vr.2) (Veltri *et al*., 2018) and iAMPpred (Meher *et al*., 2017). To obtain an unbiased estimation, we first excluded the repeated data in the training dataset of the particular predictor and the external testing dataset we prepared. The performance of each prediction model is demonstrated in Table 1. The results show that our prediction model is superior to others in terms of performance. Detailed discussion and result tables of this research are documented in Supplement 4.

**Table 1.** Comparison of our model with other predictors.

| **predictors** | accuracy | precision | sensitivity | specificity | F1 score | mcc |
| --- | --- | --- | --- | --- | --- | --- |
| APS vr.2 | 0.8097 | 0.8796 | 0.7717 | 0.8601 | 0.8222 | 0.6256 |
| iAMPpred | 0.7367 | 0.7436 | 0.7365 | 0.7368 | 0.7400 | 0.4733 |
| Our model | 0.8850 | 0.9035 | 0.8620 | 0.9080 | 0.8822 | 0.7707 |

## 4 Conclusion

We presented a web-server AMP predictor AI4AMP and a physicochemical properties-based protein-encoding method PC6. PC6 may be implemented on similar tasks that require protein-encoding. AI4AMP could serve as a beneficial tool for drug developers, as they could use the tool to discern AMPs and non-AMPs. In AI4AMP, we focused on the prediction of antibacterial peptides. Updated versions of AI4AMP are expected in the future, as newly-discovered AMPs may train our prediction model. Besides, the deep learning model is available at https://github.com/LinTzuTang/AI4AMP_predictor.

