## Supplements for "AI4AMP: Sequence-based antimicrobial peptides predictor using physicochemical properties-based encoding method and deep learning"

### Supplement 1. Data gathering and preprocessing

#### 1.1 Positive data collection

Our positive dataset was obtained from four databases: APD3 (Li *et al*., 2015), LAMP (Zhao *et al.*, 2013), CAMP3 (Thomas *et al*., 2010), and DRAMP (Kang *et al*., 2019). We downloaded all anti-bacterial AMP data from the four databases and excluded AMPs with sequence length shorter than ten amino acids, as well as those containing unusual amino acids, such as B, Z, U, X, i, and n. Also, we removed duplicated AMPs and obtained 6623 positive data in total **(Supplementary Figure 2. Panel A)**. **Supplementary Figure 1** shows the length distribution of 6623 data, and obviously, most AMPs are shorter than 50 amino acids in length.

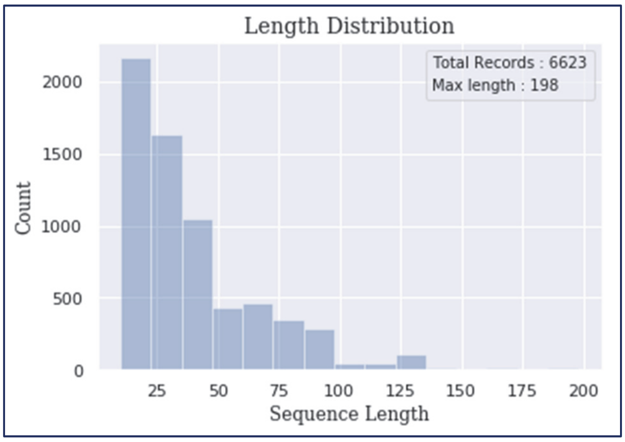


**Supplementary Figure 1.** Histogram of 6623 AMP records.

#### 1.2 Negative data collection

Our negative dataset is a combination of real-world peptides and manually generated sequences. Real-world peptides were obtained from UniProt (https://www.uniprot.org/) with the following criteria as follows. We obtained reviewed data with sequence length between 10 to 50 then excluded those without AMP-related keywords, such as 'Antimicrobial,' 'Antibiotic,' 'Amphibian defense peptide,' and 'Antiviral protein.' Manually generated sequences were randomly generated (20 essential amino acids occur in equal probabilities), with the same length distribution as the positive dataset. Finally, we obtained 6623 negative data **(Supplementary Figure 2. Panel B)**.

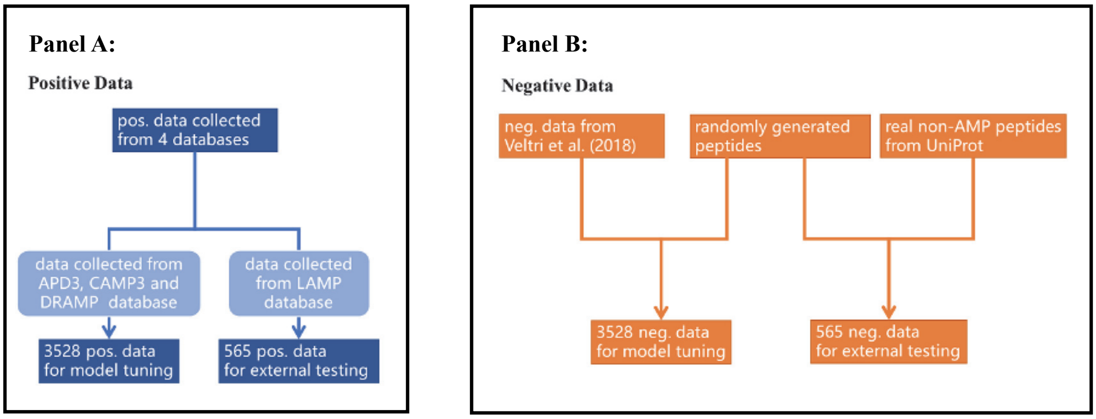


**Supplementary Figure 2.** The workflow of data collection. Positive data collection (Panel A). Negative data collection (Panel B).

#### 1.3 Data for model tuning

We used 3528 positive data and 3528 negative data to construct and tune our model. Three thousand five hundred twenty-eight positive data were obtained from 6623 positive data: after excluding the data from the LAMP dataset, we filtered out the remaining AMPs sharing above 90% sequence identity with existed AMPs (CD-HIT (Niu et al., 2010) was applied for sequence identity calculation). As for the 3528 negative data, 1778 data of which were from the existing dataset built by Veltri et al. (2018). To get an equal amount of data in the negative dataset and positive dataset, we randomly generated 1750 data with the same peptide length distribution as the positive dataset and obtained 3528 negative data. The total 7056 data (3528 negative data + 3528 positive data) were split twice with a 90:10 ratio. First, we achieved 6350 data (90%) and 706 data (10%). Seven hundred six data were used as our testing dataset, and 6350 data were further divided into 5715 data (90%) as our training dataset and 635 data (10%) as our validation dataset.

#### 1.4 Data for external testing

The purpose of distinguishing an external testing dataset from the data we used for model tuning is to challenge the ability of AMP predictors. In our external testing dataset, the data were from the LAMP database, which is distinct from the databases in which our data in the model tuning process were from. This approach could simulate real-world conditions, as peptide sequences from the different databases might have different properties, which could lower the accuracy of the prediction. An external testing dataset could tell us if our model can still reach great performance with data coming from different databases. Additionally, such an external dataset was also used for the comparison of our model to other state-of-art AMP predictors. To obtain an unbiased comparison, we excluded data if it is identical to the training dataset of each state-of-art predictor. In this way, we can observe how the AMP predictors perform with data they have never learned. The external positive testing dataset was collected from the

LAMP database, and AMPs sharing above 90% sequence identity with existed AMPs were filtered out. Finally, 565 data were obtained in the external positive testing dataset. Two hundred eighty-five of our external negative testing datasets were randomly selected from data collected from UniProt, and 280 data were randomly chosen from the arbitrarily generated sequence dataset with the same peptide length distribution as an external positive dataset. Finally, 565 data were obtained in the external negative testing dataset. In total, there are 1130 data in the external testing dataset.

##### 1.5 Data for the final model

After we confirmed the best model architecture, hyperparameters, and protein-encoding methods, we trained the model with all available data (including 6623 positive data and 6623 negative data). Please find all the datasets used in this study from our online HELP page (https://symbiosis.iis.sinica.edu.tw/PC_6/helppage.html). In the future, both of the positive and negative datasets will be continuously updated with the same criteria when newly-discovered AMPs are available.

### Supplement 2. PC6 protein-encoding method

##### 2.1 Physicochemical properties clustering analysis

Here, we collected the physicochemical properties of amino acids from the R package 'Peptides.' After that, we filtered out properties that contain "NA" in the dataset and obtained the remaining 115 properties. Then, we used the R function to calculate the correlation between each property and applied clustering analysis through hierarchical clustering. Finally, taking the K-means approach, we determined six as the optimal number of clusters (**Supplementary Figure 3**). We chose one physicochemical property for protein-encoding from each cluster named as Physiochemical Component 6 (PC6). The selected physicochemical properties are hydrophobicity (H1), the volume of side chains (V), polarity (P1), pH at the isoelectric point (pl), the negative of the logarithm of the dissociation constant for the -COOH group (pKa), and net charge index of the side chain (NCI).

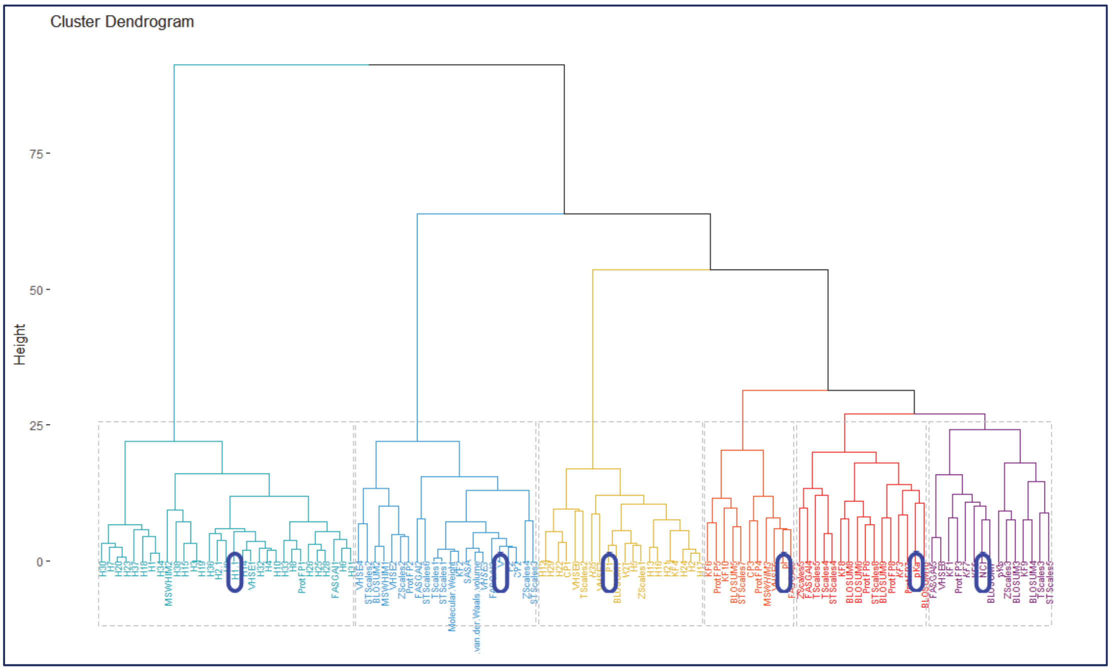


**Supplementary Figure 3.** Hierarchical clustering plot of physicochemical properties. Six selected physicochemical properties are marked in blue.

#### 2.2 Protein encoding method

An ideal protein-encoding method should take multiple protein features into consideration, which may include the order, the pattern, or the physicochemical properties of the amino acids within the protein sequence. It was discovered that the implementation of different protein-encoding methods profoundly affects the performance of a prediction model. The following studies proposed AMP prediction models with different protein-encoding and model training methods. In AmPEP (Bhadra et al., 2018), AMPs were encoded regarding the distribution patterns of amino acids, and the prediction model was built by random forest (RF). Another study, iAMPpred (Meher et al., 2017), encoded AMP considering the composition, along with physicochemical and structure properties, of amino acid, pseudo amino acid, and normalized amino acid. Besides, they constructed their model based on support vector machine (SVM). Antimicrobial Pep-tides Scanner vr.2 (Veltri et al., 2018), encoding AMPs through a neural network embedding layer, utilized deep learning methods to develop their AMP classifier. In this paper, we proposed PC6, a novel protein-encoding method. The core idea of the PC6 encoding method is to apply word embeddings regarding the physicochemical properties of each amino acid (**Supplementary Figure 4**). Each amino acid character in a sequence would be replaced by a vector composed of six physicochemical property values. We first obtained a table with 20 amino acids with its corresponding physicochemical properties. Then, we normalized the values in each physicochemical property on the same scale. The character "X" was added for sequence padding, and its corresponding values were set to 0 for all six physicochemical properties. Therefore, we generated a protein-encoding table containing 21 tokens (20 amino acids and one padding character). Considering that the sequences in our AMP dataset have a maximum length of 198, we padded all-AMPs to 200 in length. After that, we replaced each token of a sequence with six values based on the PC6 protein-encoding table and formed a 200×6 matrix. Finally, all training data would be encoded by this method to generate the input for model training.

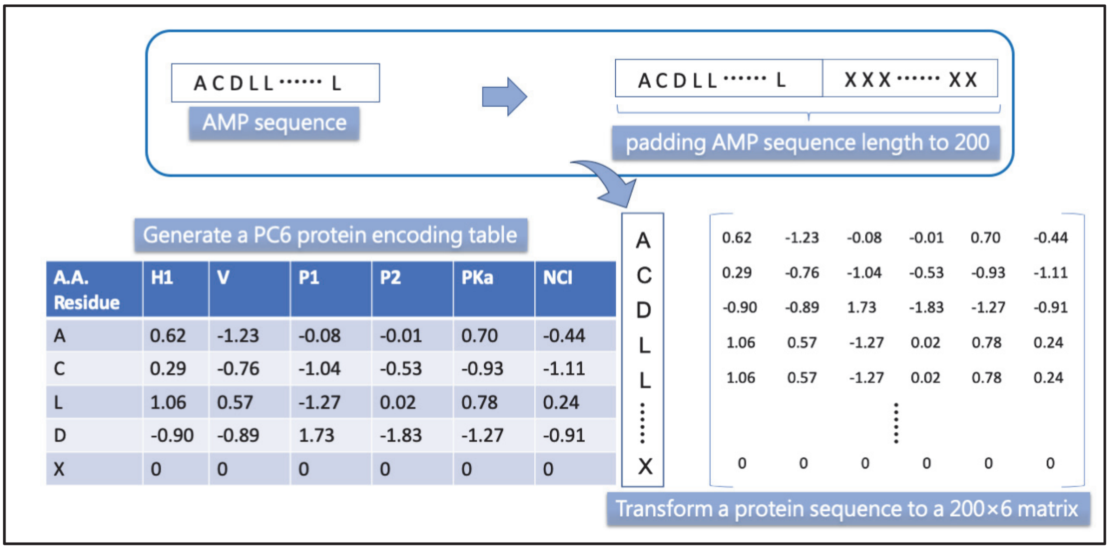


**Supplementary Figure 4.** PC6 protein-encoding method. A padded AMP will be transformed into a 200×6 matrix.

### Supplement 3. Deep neural network model

We implemented our neural network by Keras. The model architecture consists of one convolution layer, one long short-term memory (LSTM) layer, and one dense layer (**Supplementary Figure 5**), which is a typical architecture in natural language processing (NLP) tasks. We did not apply the pooling layer in our model, as it removes common features, which may result in the disappearance of some protein fragment information, according to Hassanzadeh & Wang *et al.* (2016). Additionally, only one convolution layer was used because adding more convolution layers decreased the validation accuracy in our case. This result showed consistency with that of Sun et al. (2017) 's protein-protein interaction model training. Our output layer is composed of a one-dimensional dense layer with a sigmoid activation function, which produces a value ranging from 0 to 1 (representing the probability that the peptide is an AMP). Our convolution layer was built with 64 one-dimensional filters of length 16, ReLU activation function, and LSTM layer containing 100 units. Binary cross entropy was implemented as the loss function. Adam optimizer, with a learning rate of 0.0003, was applied as our optimizer. We tuned our model by training the model with the training dataset (90%) and evaluating the performance by the validation dataset (10%). Grid search was employed for model tuning, including the adjustment of the learning rate, batch size, and optimizer. After we selected the parameters, they were unchanged throughout the process of protein-encoding comparison and model comparison. Finally, all available data (including 6623 positive data and 6623 negative data) were used to train the final model on our AI4AMP website. The final model was trained with 200 epochs, and the batch size was set to half the number of the training dataset. To avoid overfitting, early stopping was performed.

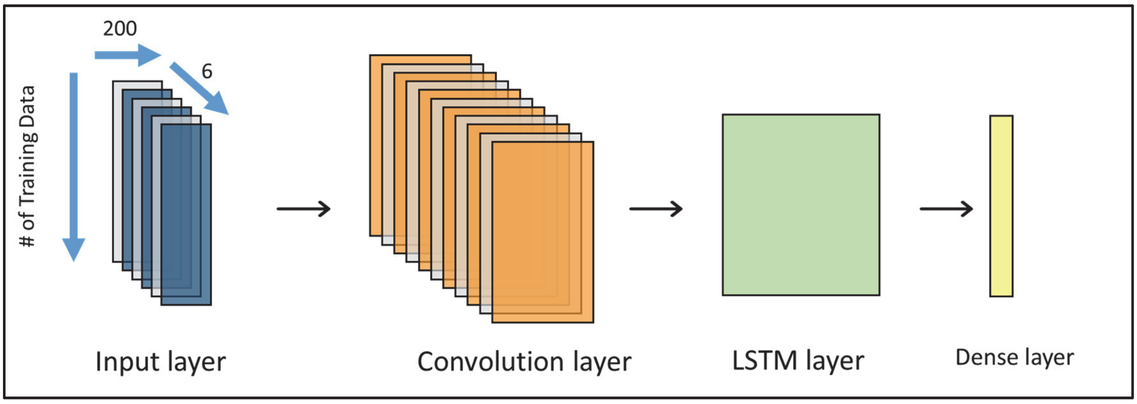


**Supplementary Figure 5.** Our model architecture. After PC6 encoding, protein sequences will pass through one convolution layer, one long short-term memory (LSTM) layer, and one dense layer.

### Supplement 4. Discussion and Result

#### 4.1 Comparison of protein-encoding methods

In order to compare the four protein-encoding methods, we tested the performance of the same deep learning model (Supplement 3) with four different protein-encoding methods, including the autocovariance method (AC), embedding layer, Word2Vec, and PC, respectively, as shown as **Supplementary Table 1**. The AC7 encoding methods were derived from seven physicochemical properties used in Sun et al. (2017). Moreover, we developed AC6 based on AC7 by replacing six physicochemical properties mentioned in supplementary figure 3. As the same concept, we also adapt those seven physicochemical properties used in AC7 into PC7. Although the AC encoding method had an exceptional performance in protein-protein interaction prediction, we found out that it did not fit well in our AMPs prediction task. The AC encoding method was far behind the other encoding methods in terms of accuracy. We suppose that too much sequence information was lost after encoding because AMPs are generally short in length (20-50 amino acids on average) and that the lag parameter value in the AC equation was set to 10, a relatively small number. Moreover, if we tested the models by an external dataset, the results indicated that models encoded with six physicochemical properties performed better than seven physicochemical properties (**Supplementary Table 2**). Except for the AC encoding method, the other three encoding methods achieved outstanding performances in terms of accuracy (ACC) and Matthews correlation coefficient (MCC). Therefore, we then evaluated the stability of the three encoding methods through 10-fold validation, as shown in **Supplementary Table 3**. PC and Word2Vec encoding methods achieved higher accuracy than the embedding layer encoding method.

**Supplementary Table 1.** Comparison of protein-encoding methods in the same setting (testing dataset).

| Encoding Method | accuracy | precision | sensitivity | specificity | F1 score | MCC |
|---|---|---|---|---|---|---|
| AC7 | 0.7507 | 0.7580 | 0.7365 | 0.7649 | 0.7471 | 0.5016 |
| AC6 | 0.7040 | 0.7130 | 0.6827 | 0.7252 | 0.6975 | 0.4083 |
| embedding layer | **0.8952** | 0.9091 | **0.8782** | 0.9122 | 0.8934 | **0.7908** |
| Word2Vec | 0.9065 | **0.9415** | 0.8669 | **0.9462** | **0.9027** | 0.8156 |
| PC7 | 0.8782 | 0.9083 | 0.8414 | 0.9150 | 0.8735 | 0.7584 |
| PC6 | 0.8895 | 0.9205 | 0.8527 | 0.9263 | 0.8853 | 0.7812 |

**Supplementary Table 2.** Comparison of protein-encoding methods in the same setting (external testing dataset).

| Encoding Method | accuracy | precision | sensitivity | specificity | F1 score | MCC |
|---|---|---|---|---|---|---|
| AC7 | 0.7311 | 0.8152 | 0.6248 | 0.8464 | 0.7074 | 0.4808 |
| AC6 | 0.7818 | 0.8534 | 0.7009 | 0.8695 | 0.7697 | 0.5760 |
| embedding layer | 0.8690 | **0.9096** | 0.8195 | **0.9186** | 0.8622 | 0.7417 |
| Word2Vec | 0.8460 | 0.8574 | 0.8301 | 0.8619 | 0.8435 | 0.6924 |
| PC7 | 0.8610 | 0.8849 | 0.8301 | 0.8920 | 0.8566 | 0.7235 |
| PC6 | **0.8850** | 0.9035 | **0.8620** | 0.9080 | **0.8822** | **0.7707** |

**Supplementary Table 3.** Models stability of three encoding methods evaluation through 10-fold validation.

| | Embedding layer | PC6 | Word2Vec |
|---|---|---|---|
| Average accuracy | 0.8458 | 0.8749 | **0.8838** |

#### 4.2 Testing of traditional machine learning methods

To compare the performance of deep learning and traditional machine learning. We applied PC6 and Word2Vec encoding methods, respectively, along with two machine learning models: a support vector machine (SVM) model and a random forest model. The performance of each combination was shown in **Supplementary Table 4.** Both PC6 and Word2Vec models trained by deep learning performed better than models trained by SVM and random forest. We, therefore, concluded that a deep learning approach is necessary for our task, as it may learn the information of AMPs more exhaustively, compared to the traditional machine learning approach. Otherwise, we found that the encoding method adopting PC6 was better than Word2Vec with any machine learning algorithm when evaluated using the external testing dataset (**Supplementary Table 5**).

**Supplementary Table 4.** Comparison of models trained by other machine learning algorithms (testing dataset).

| Encode / Machine learning | accuracy | precision | sensitivity | specificity | F1 score | MCC |
|---|---|---|---|---|---|---|
| PC6 / SVM | 0.8329 | 0.8550 | 0.8017 | 0.8640 | 0.8275 | 0.6670 |
| PC6 / random forest | 0.8385 | 0.8770 | 0.7875 | 0.8895 | 0.8299 | 0.6806 |
| PC6 / deep learning | 0.8895 | 0.9205 | 0.8527 | 0.9263 | 0.8853 | 0.7812 |
| Word2Vec / SVM | 0.8300 | 0.8499 | 0.8017 | 0.8584 | 0.8251 | 0.6611 |
| Word2Vec / random forest | 0.8499 | 0.8800 | 0.8102 | 0.8895 | 0.8437 | 0.7019 |
| Word2Vec / deep learning | **0.9065** | **0.9415** | **0.8669** | **0.9462** | **0.9027** | **0.8156** |

**Supplementary Table 5.** Comparison of models trained by other machine learning algorithms (external testing dataset).

| Encode / Machine learning | accuracy | precision | sensitivity | specificity | F1 score | MCC |
|---|---|---|---|---|---|---|
| PC6 / SVM | 0.8566 | 0.8592 | 0.8531 | 0.8602 | 0.8561 | 0.7133 |
| PC6 / random forest | 0.8513 | 0.8869 | 0.8053 | 0.8973 | 0.8442 | 0.7057 |
| **PC6 / deep learning** | **0.8850** | **0.9035** | **0.8620** | **0.9080** | **0.8822** | **0.7707** |
| Word2Vec / SVM | 0.7327 | 0.7514 | 0.6956 | 0.7699 | 0.7224 | 0.4668 |
| Word2Vec / random forest | 0.7858 | 0.8549 | 0.6885 | 0.8832 | 0.7627 | 0.5828 |
| Word2Vec / deep learning | 0.8460 | 0.8574 | 0.8301 | 0.8619 | 0.8435 | 0.6924 |

##### 4.3 Comparison of our models with other state-of-art predictors

We performed a comparison of our deep learning models, encoded by PC6 and Word2Vec methods separately, with other state-of-art AMP predictors. An external testing dataset tested the performance of each predictor. As shown in **Supplementary Table 6**, the PC6 model had the best external test performance. We also demonstrated the number of misclassified cases (external testing dataset) of AI4AMP(our final model using PC6 encoding method) and Antimicrobial Peptide Scanner vr.2 in **Supplementary Figure 6.** We observed significantly better performance of AI4AMP due to less false-positive cases. It was obvious that AI4AMP was more capable of discerning negative data. We listed the overlapping misclassified cases of AI4AMP and Antimicrobial Peptide Scanner in **Supplementary Table 7, 8**, as it would be interesting to investigate the sequences that are difficult to recognize for both models.

**Supplementary Table 6.** Comparison of AMP predictors.

| predictors | accuracy | precision | sensitivity | specificity | F1 score | MCC |
|---|---|---|---|---|---|---|
| Antimicrobial Peptide Scanner vr.2 * | 0.8097 | 0.8796 | 0.7717 | 0.8601 | 0.8222 | 0.6256 |
| iAMPpred ** | 0.7367 | 0.7436 | 0.7365 | 0.7368 | 0.7400 | 0.4733 |
| **PC6** | **0.8850** | **0.9035** | **0.8620** | **0.9080** | **0.8822** | **0.7707** |

*: https://www.dveltri.com/ascan/v2/

**: http://cabgrid.res.in:8080/amppred/

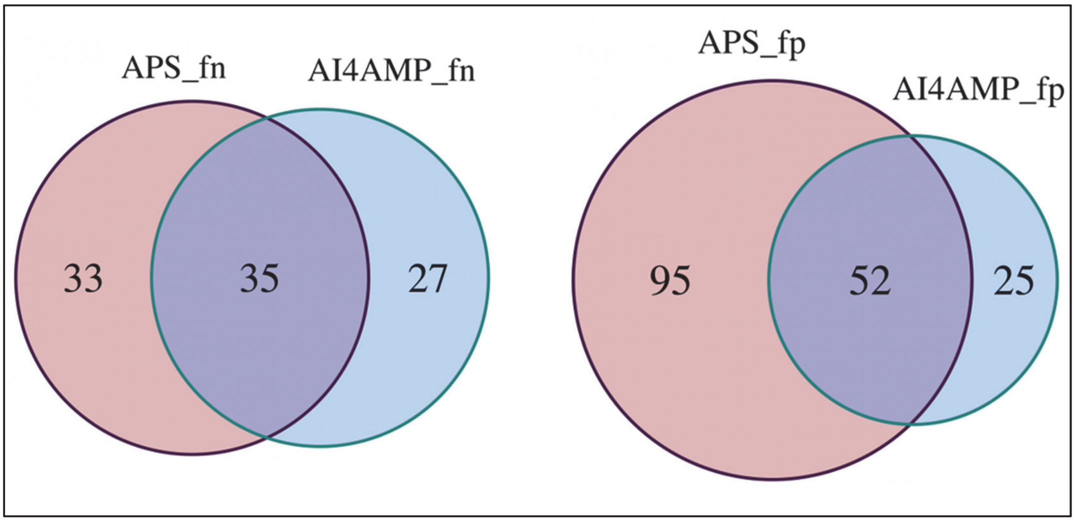


**Supplementary Figure 6.** The Venn diagram presents the AMP prediction overlaps and differences between AI4AMP (our final model) and Antimicrobial Peptide Scanner vr.2. The overlaps and differences of **false-negative** prediction of the models (Left). The overlaps and differences of **false-positive** prediction of the models (Right).

**Supplementary Table 7.** A list of overlapping false-negative cases.

| | seq_name | sequence |
|---|---|---|
| 1 | L01A001278 | ADSGEGDFLAEGGGVR |
| 2 | L01A001277 | ADSGEGDFLAEGGGVRKLIK |
| 3 | L01A001271 | ALYKKKIIKKLLES |
| 4 | L01A001289 | ATKKNGRKLCLDLQAAL |
| 5 | L01A001979 | DYQAKLAAYQKEL |
| 6 | L01A001550 | EAQTRCQVAGWGSQSRSGGR |
| 7 | L01A001276 | EGVNDNEEGFFSA |
| 8 | L01A001280 | EWVQKYVSDLELSAWKKILK |
| 9 | L01A001955 | GSSKSPSKDKDKDPGDC |
| 10 | L01A002130 | IEDIAVNSKYQGQGLGKLLIPRT |
| 11 | L01A002128 | IEDISVAKSEQGKKLGYYLV |
| 12 | L01A001777 | IGEDVYTPGISGDSLR |
| 13 | L01A001281 | KWVREYINSLEMSKKGLAG |
| 14 | L01A002134 | LEDFFVMSDYRGFGIGSEIL |
| 15 | L01A002138 | LFHLSVDNEHRGQGIAKALV |
| 16 | L07APD0074 | MVPTTFTLTTNNFLSDYQQLFI |
| 17 | L01A002135 | NEPSINFYKRRGA |
| 18 | L01A002139 | QLSAMGLYQSLGF |
| 19 | L01A001837 | RERDHELRHRRHHHQ |
| 20 | L01A001619 | RLVERIRQLTASRQLIPQLIQYV |
| 21 | L01A000716 | RSTEDIIKSISGGGFLNAMNA |
| 22 | L01A001287 | SAIHPSSILKLEVICIGVLQ |
| 23 | L01A001959 | SPSNETPKKKKKRFSFKKSG |
| 24 | L01A001387 | SSSKEENRIIPGGI |
| 25 | L01A001279 | SWVQEYVYDLEL |
| 26 | L01A002132 | VEDVVVSDECRGKQLGKLLL |
| 27 | L01A001286 | YAERLCTCSIKAEV |
| 28 | L01A001597 | YHELRDLLLIVTRIVELLGRE |
| 29 | L01A001644 | YHRLLRDLLIVTRIVELL |
| 30 | L01A001651 | YHRLLRDLLIVTRIVELLGRR |
| 31 | L01A001641 | YHRLRDLALIVTRIVELL |
| 32 | L01A001643 | YHRLRDLLLIVTRIVELL |
| 33 | L01A001362 | YQVIQSWEHWRE |
| 34 | L01A002136 | YSTGMVHLLLQVTIDGRNYI |
| 35 | L102675000 | GSAQPYKQLHKVVNWDPYG |

**Supplementary Table 8.** A list of overlapping false-positive cases.

| | seq_name | sequence |
|---|---|---|
| 1 | non_amp_4 | MGGVGKTKRMRRLKRKRRKMRQRSK |
| 2 | non_amp_19 | INIKDILAKLVKVLGHV |
| 3 | non_amp_36 | MDIVSLAWAALMVVFTFSLSLVVWGRSGL |
| 4 | non_amp_37 | FLPMIAKLLGGLL |
| 5 | non_amp_46 | INWKAIIEAAKQAL |
| 6 | non_amp_47 | METFCYMKWPVRHHKSRRVSH |
| 7 | non_amp_49 | RGCPRILMRCKRDSDCLAGCVCQKNGYCG |
| 8 | non_amp_53 | MDIVSLAWAALMVVFTFSLSLVVWGRSGL |
| 9 | non_amp_61 | PVQSVFSATRCGAN |
| 10 | non_amp_69 | MVTMVKKWLLLMTFMAGCRGMIY |
| 11 | non_amp_74 | MDIVSLAWAALMVVFTFSLSLVVWGRSGL |
| 12 | non_amp_83 | FLFFAFPHPL |
| 13 | non_amp_112 | MRAKWKKKRMRRLKRKRRKMRQRSK |
| 14 | non_amp_115 | MDIVSLAWAALMVVFTFSLSLVVWGRSGL |
| 15 | non_amp_151 | MVKLRLKRCGRK |
| 16 | non_amp_175 | GFKNVALSTARGF |
| 17 | non_amp_181 | MDIVSLAWAALMVVFTFSLSLVVWGRSGL |
| 18 | non_amp_187 | MNAAIFRFFFYFST |
| 19 | non_amp_192 | MRAKWRKKRVRRLKRKRRKVRARSK |
| 20 | non_amp_198 | GAPICGESCFTGKCYTVQCSCSWPVCTRN |
| 21 | non_amp_204 | GLPVCGETCFGGRCNTPGCTCSYPICTRN |
| 22 | non_amp_214 | MDIVSLAWAALMVVFTFSLSLVVWGRSGL |
| 23 | non_amp_227 | GIGSALANAAKLVAGIV |
| 24 | non_amp_228 | ARRRRRSSRPQRRRRRRRHGRRRRGRR |
| 25 | non_amp_229 | GLPVCGETCFGGTCNTPGCSCSYPICTRN |
| 26 | non_amp_231 | INWLKLGKAVIDAL |
| 27 | non_amp_233 | MKKARRSPSRRKGARLWYVGGSQF |
| 28 | non_amp_240 | MDIVSLAWAALMVVFTFSLSLVVWGRSGL |
| 29 | non_amp_243 | MRAKWRKKRMRRLKRKRRKMRQRSK |
| 30 | non_amp_270 | ARVPFYRYYK |
| 31 | non_amp_332 | KWQSFPACWICWI |
| 32 | non_amp_348 | GLIGRALKYISTRCMTHTDD |
| 33 | non_amp_352 | NLFENWLSCFEYRMLRNWCRMTD |
| 34 | non_amp_368 | NYKQPFFMNIWKTHRFHV |

| | | |
|---|---|---|
| 35 | non_amp_390 | TLPIRLRKYCEWAHACPMLG |
| 36 | non_amp_411 | EGWPYIWGVHSFVCNQCKEYI |
| 37 | non_amp_415 | TVQVVGFRWSHKWFRFQKWW |
| 38 | non_amp_417 | VTTEWQAIKRYMMKHWAKH |
| 39 | non_amp_454 | KSWYYTAVWWLGVCNTWPFAFQLS |
| 40 | non_amp_467 | PAFCQCRLWCRHIKQMNWMS |
| 41 | non_amp_469 | VNKLHQSMMQCWNCACHMA |
| 42 | non_amp_473 | YRDGWVILYKGCKLGGHR |
| 43 | non_amp_481 | PRTRWYHHKYMIWRLLTAGWIHSL |
| 44 | non_amp_483 | THHHAPWGCCTYHL |
| 45 | non_amp_486 | PFWNSWFYQTASVKIRGYQG |
| 46 | non_amp_505 | HHMCHTACAWGGWHTF |
| 47 | non_amp_522 | WWPCWAMNCKFWC |
| 48 | non_amp_534 | KWKFDGPVGYDGLCPPAFVT |
| 49 | non_amp_541 | AKTWGKQLERLDCMKTITAFYSCRWVKG |
| 50 | non_amp_562 | FYGFWIEKIKQYLT |
| 51 | non_amp_564 | THHIPLGPCCFY |
| 52 | non_amp_565 | MYWPLLPCFQNFICAPWHEV |

#### 4.4 Evaluation of our PC6 model and Antimicrobial Peptide Scanner vr.2

To ensure it is PC6 protein-encoding method and our model architecture that result in superior performance, not just the fact that more training data was used. Here, we trained our model with the same dataset (2,021 AMP and Non-AMP sequences, respectively) used in Antimicrobial Peptide Scanner vr.2 (vr.2 Feb2020). Then, we used our external testing dataset to evaluate the performance of our model and the model provided by Antimicrobial Peptide Scanner vr.2 (the latest version: Feb2020 model). Overall, our PC6 model had better performance, as shown in **Supplementary Table 9.**

**Supplementary Table 9.** Comparison of PC6 model and Antimicrobial Peptide Scanner vr.2 (external testing dataset).

| predictors | accuracy | precision | sensitivity | specificity | F1 score | MCC |
|---|---|---|---|---|---|---|
| Antimicrobial Peptide Scanner vr.2 * | 0.8097 | **0.8796** | 0.7717 | 0.8601 | 0.8222 | 0.6256 |
| **PC6** | **0.8619** | 0.8619 | **0.8619** | **0.8619** | **0.8619** | **0.8619** |
